# Role of Early-Life Microbiome Colonization in Physiological Development of Drosophila melanogaster

**DOI:** 10.64898/2026.08.16.745128

**Authors:** Zihan Tian, William B. Ludington

## Abstract

The influences of the gut microbiome on animal physiology are well-documented, yet the developmental timing of microbial colonization and its long-term consequences remain poorly understood. In this study, we investigated how the timing of bacterial colonization during development affects transcriptional programming and phenotypic outcomes in adult *Drosophila melanogaster* reared on a common, rich diet. Using RNA-seq analysis on whole flies colonized either as newly hatched larvae or as newly eclosed adults, we observed minor but distinct transcriptional responses dependent on when flies were colonized. Both embryonic and adult colonization were associated with ~ 25 to ~ 200 differentially expressed genes compared to axenic controls, with the majority upregulated and enriched for immune-response genes, suggesting that colonization establishes a broader immune competence. However, only 10 genes showed persistent differential expression that was not normalized by introducing bacteria to adult axenic flies, including mitochondrial genes, the adipokinetic hormone (*Adh*), and a putative secreted neuropeptide. Overall, these findings suggest that *Drosophila* development on a rich diet is largely robust to the timing of bacterial colonization but that certain metabolic effects may occur.

## Introduction

The composition and abundance of intestinal microbial communities have been shown to modulate diverse aspects of host biology, including nutrient acquisition, immune system development, metabolic homeostasis, and even neurological function (Grenier and Leulier, 2020; Lesperance and Broderick, 2020; Trinder et al., 2017). Dysbiosis, or imbalance in microbial composition, has been associated with numerous conditions, including inflammatory bowel disease, obesity, and autoimmune disorders (Zhang et al., 2015). Understanding the mechanisms by which gut microbes shape host phenotypes has become a central question in biology, with implications for both basic science and translational medicine.

In mammals, the “hygiene hypothesis” states that reduced microbial exposure during early life could lead to increased susceptibility to various diseases, including autoimmune diseases and cancer (Dujon et al., 2023). This concept has been extended to the developmental origins of health and disease framework, which proposes that early-life environmental exposures, including microbial colonization, can have lasting effects on adult health outcomes. In the context of allergic diseases, the gut microbiome has emerged as a critical early-life factor that is capable of influencing the risk of chronic disease (Sbihi et al., 2019). Lacking early exposure to infectious agents influences risk for immune-mediated diseases, including asthma and allergies (Al Nabhani and Eberl, 2020; Gensollen et al., 2016; Sbihi et al., 2019). This concept of a critical developmental window suggests that the timing of initial microbial colonization could have long-term physiological effects (Metcalf et al., 2022; Warne et al., 2019).

Due to several key advantages, such as a simplified microbiome with low complexity and diversity, *Drosophila melanogaster* is a powerful model organism for investigating host-microbiome interactions (Ludington and Ja, 2020; Trinder et al., 2017). Gut-associated microbes of the fruit fly are relatively simple, typically comprising 5-20 bacterial species in laboratory-reared animals, dominated by members of the Lactobacillaceae and Acetobacteraceae families (Ludington et al., 2025; Miguel-Aliaga et al., 2018). Compared to the hundreds of species found in mammalian guts, this simplified microbial community facilitates mechanistic studies of specific host-microbe interactions. In a laboratory setting, *Drosophila* embryos can be readily rendered germ-free (axenic) through surface sterilization with dilute bleach, and subsequently maintained germ-free for hundreds of generations (Northrop, 1926). These germ-free flies can then be colonized with defined microbial communities to generate strains of gnotobiotic animals, allowing precise manipulation of microbial exposure timing and composition (Kietz et al., 2018; Koyle et al., 2016).

The gut microbiota composition exerts pronounced effects on *Drosophila* development when deprived of standard nutrition, with germ-free larvae exhibiting delayed development, reduced body size, and altered metabolic profiles compared to conventionally-reared counterparts(Grenier and Leulier, 2020; Matos et al., 2017; Shin et al., 2011; Storelli et al., 2018; Téfit and Leulier, 2017). However, minimal differences in development and fecundity were reported for conventionally-reared versus axenic flies on a nutritionally complete diet (Northrop, 1926). Moreover, the adult *Drosophila* gut transcriptome on a nutritionally-complete diet is largely insensitive to the identity of colonizing bacterial species, though whole-body transcriptional profiles differ between conventionally-reared and gnotobiotic flies, likely due to yeast in the conventional flies (Elya et al., 2016).

The widespread use of gnotobiotic flies is inconsistent between labs and could be a source of differences in experimental outcomes (Douglas, 2018). For instance, some labs generate germ-free and gnotobiotic flies by bleaching the embryos, which is known to reduce lifespan (Lee et al., 2019), before rearing under aseptic or gnotobiotic conditions. Others treat newly eclosed adults with antibiotics to kill off bacteria that survived metamorphosis (Fast et al., 2018; Ryu et al., 2008; Sansone et al., 2015). Still others maintain germ-free stocks in perpetuity, which has been shown since 1926 to have minimal effects on fly health and development when reared on a nutritionally replete diet (Northrop, 1926). In this study, we investigate the impact of the timing of bacterial colonization on *D. melanogaster* physiology by comparing three experimental groups: (1) flies colonized with a defined seven-species bacterial community (Eble et al., 2023; Yang et al., 2025), (2) germ-free flies that receive bacterial colonization only upon adult eclosion, and (3) flies maintained germ-free. We assess physiological outcomes through measurements of body size (length and mass) and whole-transcriptome mRNA sequencing (RNAseq) at eclosion and 1 week post-eclosion.

## Results

### Bacterial colonization has minor effects on adult body size

We first developed an experimental strategy to measure the impacts of the timing of bacterial colonization on fly physiology (Fig. 1), with five treatment groups. We started with a CantonS, Wolbachia-free fly stock that has been maintained germ-free for ≈10 generations to avoid any detrimental effects of bleaching, which have been observed in the first generation after treatment (Lee et al., 2019). We then inoculated one set of flies with a 7-bacterial species consortium of commensal lactobacilli and acetobacters that were isolated from a single wild-caught *D. melanogaster* (Yang et al., 2025), keeping another set germ-free, yielding two parallel treatment groups. The colonized flies were maintained with bacteria for ≥ 2 generations. Flies were checked for bacteria 10 days post-eclosion. Axenic flies remained germ-free. Colonized flies maintained their full complement of bacterial morphologies and were reinoculated at each generation to ensure full exposure.

**Figure 1:**
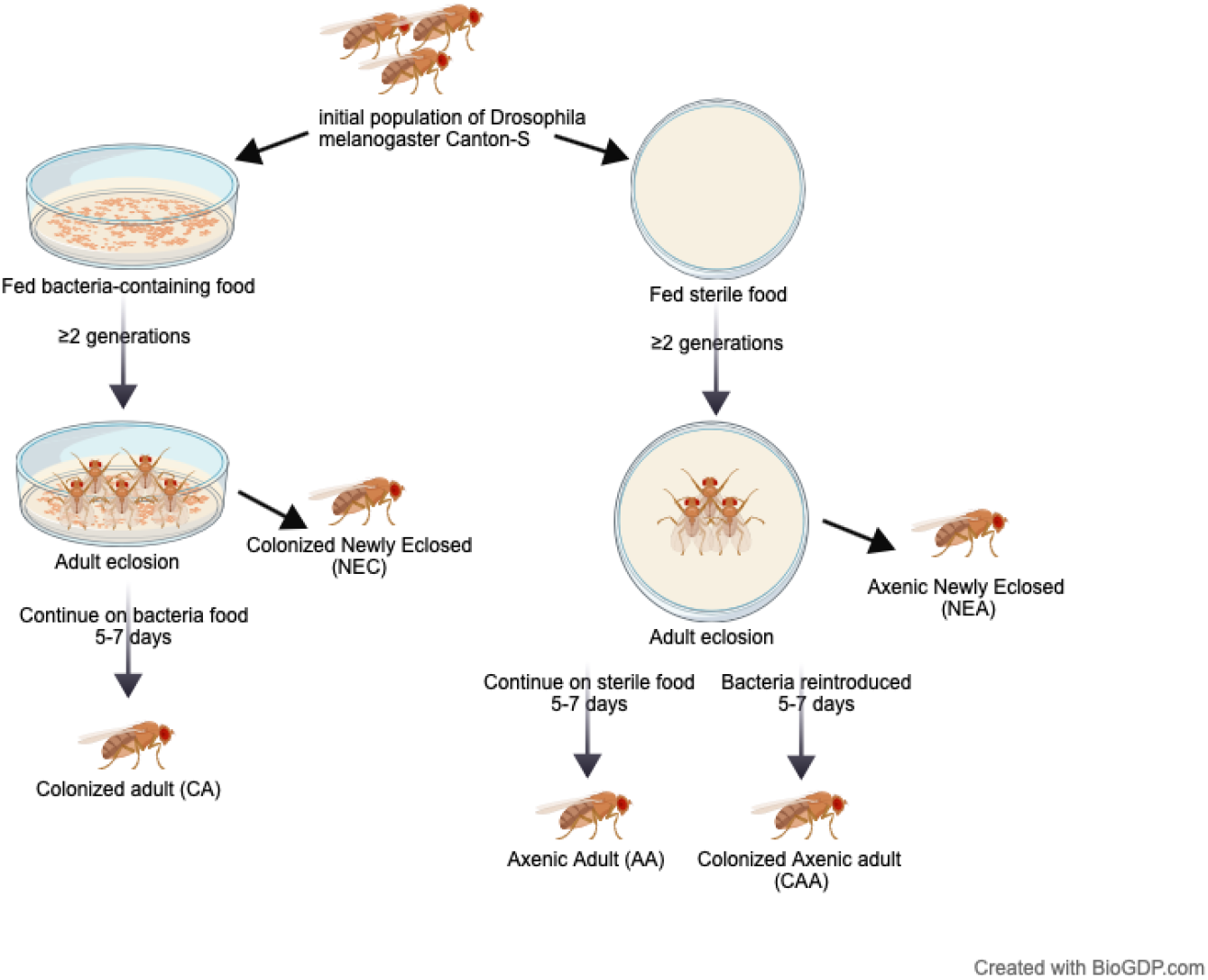
Experimental design.

We sampled the flies at eclosion for body size and RNAseq, yielding the first developmental time-point sample of the treatment groups, newly-eclosed colonized flies (NEC) and newly-eclosed axenic (NEA).

At eclosion, one group of axenic flies was transferred to food inoculated with the 7-bacteria consortium, yielding a 3rd treatment. At 6 days post-eclosion, we sampled the flies from the three treatments for body size and RNAseq, yielding colonized adult (CA), axenic adult (AA), and colonized axenic adult (CAA).

Body mass and body length were measured across experimental groups to assess the phenotypic effects of bacterial colonization timing (Figure S1). As expected, we observed significant differences between males and females (sexual dimorphism) and between newly eclosed and mature adults (developmental growth), consistent with well-established patterns in *Drosophila* biology. However, there was not a noticeable effect of bacterial inoculation.

### Bacterial colonization determines the magnitude and nature of transcriptional responses

To establish whether there were physiological differences not apparent by body size, the whole body transcriptome was measured for each of the five treatment groups. Differential gene expression analysis was performed using RNAseq and the DESeq2 (v1.42.0) analysis package in R (Love et al., 2014). The reads were fit to a gene model with 12,482 transcribed regions of the *D. melanogaster* reference genome (BDGP6.54, Ensembl release 115) using the nf-core/rnaseq pipeline (Ewels et al., 2020) across 57 samples (Table 1).

**Table 1:** Sample size (*n*) for each experimental group after filtering.

| Group | Description | n |
| --- | --- | --- |
| Colonized adult (CA) | Embryonic inoculation, 5–7 d post-eclosion | 10 |
| Colonized newly eclosed (NEC) | Embryonic inoculation, right after eclosion | 14 |
| Axenic newly eclosed (NEA) | Axenic control, right after eclosion | 11 |
| Colonized Axenic adult (CAA) | Adult recolonization, 5–7 d post-eclosion | 11 |
| Axenic adult (AA) | Axenic control, 5–7 d post-eclosion | 11 |

**Table 2:** Functional annotation of the 10 DEGs between the two colonized treatments that is not detectable against the axenic control.

| Gene Name | FlyBase ID | $\log_2FC$ | Putative Functional |
| --- | --- | --- | --- |
| snRNA:U1:21D | FBgn0003916 | 7.60 | U1 small nuclear RNA; spliceosome component |
| mt:ND4L | FBgn0013683 | 7.03 | Mitochondrial NADH dehydrogenase subunit 4L |
| mt:ND3 | FBgn0013681 | 2.23 | Mitochondrial NADH dehydrogenase subunit 3 |
| CG14075 | FBgn0036835 | 1.80 | Putative secreted peptide / putative neuropeptide |
| Akh | FBgn0004552 | 1.71 | Adipokinetic hormone; neuropeptide |
| roo{ }339 | FBti0019148 | 1.47 | roo transposable element |
| CG3699 | FBgn0040349 | 1.36 | Uncharacterized protein / putative age-associated gene |
| CG10910 | FBgn0034289 | 0.70 | Uncharacterized protein |
| Mal-A2 | FBgn0002569 | 0.52 | Maltase A2; α-glucosidase |
| CG9760 | FBgn0036259 | -1.04 | Uncharacterized protein |

**Table 3:** Target OD_600_ values for each bacterial strain to reach 10^6^ CFU/mL.

| Strain | Target OD<sub>600</sub> |
| --- | --- |
| *L. plantarum* (ZTG301) | 0.08 |
| *L. brevis* (ZTG304) | 0.30 |
| *A. malorum* (ZTG303) | 0.45 |
| *A. tropicalis* (ZTG310) | 0.12 |
| *A. orientalis* (ZTG331) | 0.30 |
| *A. cerevisiae* (ZTG313) | 0.30 |
| *A. sicerae* (ZTG360) | 0.30 |

Principal component analysis (PCA) of gene expression for all sample data revealed clustering by developmental stage, separating newly-eclosed from 6-day-old flies (Figure S2). Within the newly eclosed samples (Figure 2A), there was substantial overlap between colonized and axenic samples, which is consistent with the metamorphosis-associated gut remodeling and antimicrobial peptide surge at pupariation (Nunes et al., 2021). Past studies have found that bacterial loads are greatly reduced during pupariation (Johnston and Rolff, 2015), and our data suggest that bacterial presence during larval development had little influence on pupal development, with axenic and conventionally-reared flies emerging with an equivalent transcriptome. Consistent with this overlap, differential expression analysis between newly eclosed colonized and axenic flies (NEC vs. NEA) identified 26 DEGs (14 up-regulated and 12 downregulated), nearly an order of magnitude fewer than the 201 DEGs detected between colonized and axenic 6-day-old adults. Although no strongly enriched pathways were identified, this gene set included upregulated antimicrobial peptides (*DptA, DptB, AttC*) and the neuropeptide *Akh*, suggesting that embryonic colonization may contribute to immune and metabolic priming by the time of eclosion.

**Figure 2:**
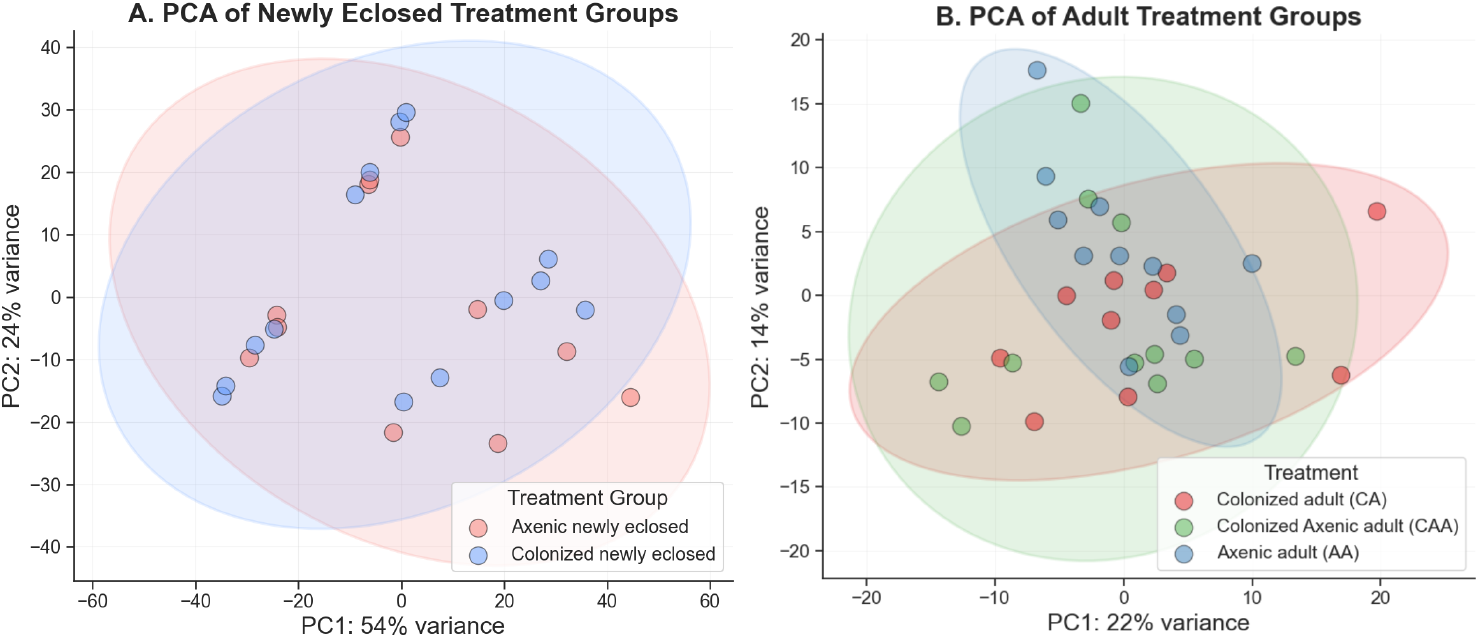
Principal Component Analysis (PCA) of RNA-seq samples. Colors indicate treatment groups as defined in the legend. **A)** Newly eclosed samples show almost complete overlap between treatment groups. **B)** The three adult treatment groups showed partial overlap in the PCA analysis. Shaded ellipses represent approximate 95% confidence regions for each group, based on the covariance matrix of PC1 and PC2 coordinates under a bivariate normal assumption. Points represent individual samples. The percentage of variance explained by each principal component is shown on the axis labels.

For adult treatment groups (Figure 2B), colonized axenic adult samples overlapped broadly with both lifelong axenic adults and colonized adults, whereas axenic adults and colonized adults showed partial separation along the principal components, suggesting distinct transcriptional profiles.

To assess individual genes that were differentially expressed between the treatment groups for each developmental stage, we used DESeq with a Benjamini–Hochberg adjusted *p*-value ≤ 0.05 and | log_2_ (fold change) | ≥0.5. We observed relatively few transcriptional changes across pairwise comparisons given the potential impacts of bacteria (Figure 3). The most extensive transcriptional reprogramming occurred in comparisons between germ-free adults and those colonized from birth, which revealed 201 DEGs (151 upregulated, 50 downregulated), consistent with previous studies (Elya et al., 2016). Comparing adults that were colonized after eclosion with germ-free adults revealed 88 DEGs (45 upregulated, 43 downregulated). Comparing adults that were colonized after eclosion with adults that were colonized for ≥ 2 generations revealed only 29 DEGs, with 25 upregulated and 4 downregulated. Overall, these results indicate a significant effect of colonization on adult gene expression.

**Figure 3:**
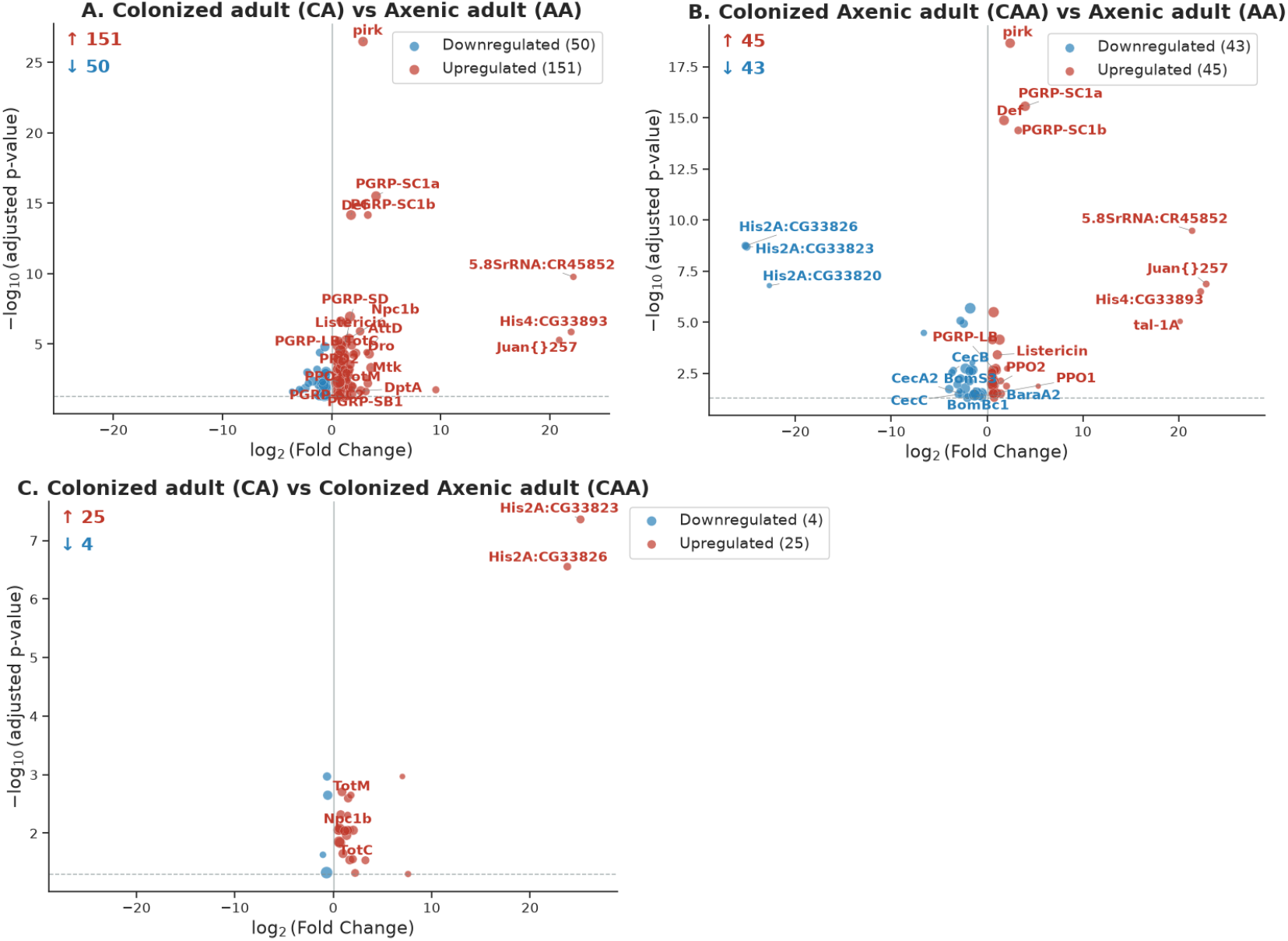
Pairwise differential gene expression of adult groups. The size of the dot represents the expression level. Labeled genes include key immune and metabolic genes of interest. Genes with extreme fold-change estimates | (log_2_ *extFC* |*>* 20) exhibited a sporadic expression pattern, where a small number of samples contained the majority of reads while the remainder had zero counts. For example, two histone H2A variants (*His2A:CG33823, His2A:CG33826*) passed the count filter (total counts more than 2,000) but had reads in only 5 of 32 samples, with zero counts in all CAA samples. This pattern is inconsistent with genuine biological regulation across replicates and likely reflects ambiguous mapping among the highly similar histone H2A paralogous genes. These genes were not interpreted biologically.

### Pathway enrichment analysis

To further examine the differentially regulated genes, we performed pathway enrichment analysis using Metascape. Overall, the pathway most affected by the timing of colonization was immune gene expression.

GO term enrichment comparing CA versus AA found major differences in (i) defense response to Gram positive bacterium, (ii) peptidoglycan metabolism, and (iii) glucuronate catabolism. To further understand the biological directionality of transcriptional changes, separate pathway enrichment analyses on upregulated and downregulated genes was performed. Colonized flies were significantly enriched for immune response against different pathogens, specifically against Gram-positive bacterium (*log*_10_*P* = − 16.70, 15 genes). Key genes included peptidoglycan recognition proteins (*PGRP-SC1a, PGRP-SC1b, PGRP-SD, PGRP-LB*) and antimicrobial peptides (*Dro, Def, Mtk, DptA, AttD, Listericin*).

Among the 50 downregulated genes, developmental processes such as muscle cell development and cell fate specification showed enrichment, consistent with tradeoffs between immune defense and somatic development (Lazzaro, 2015).

Comparing colonized axenic adults versus lifelong axenic adults, of the 88 differentially expressed genes, the most significant was the activation of the immune system process. Upregulated genes were enriched for defense response to Gram-positive bacterium (*q* = 3.8∗10^*−*6^, *log*_10_*P* = −9.37, 7 genes) and peptidoglycan catabolic process (*q* = 2.5∗10^*−*3^, *log*_10_*P* = −5.83, 3 genes), driven by pattern recognition receptors (PGRP-SC1a, *log*_2_FoldChange = 3.98; PGRP-SC1b, *log*_2_FoldChange =3.26; PGRP-LB, *log*_2_FoldChange=0.80) and select antimicrobial peptides (Listericin, *log*_2_FoldChange = 1.08). Notably, downregulated genes were significantly enriched for humoral immune response (*q* = 3.3∗10^*−*5^, *log*_10_*P* = −8.25, 6 genes), specifically the Toll and Imd signaling pathways, as shown in the overall enrichment heatmap. The downregulated effector genes included Cecropins (CecA2, *log*_2_FoldChange=−2.65; CecB, *log*_2_FoldChange=−3.51; CecC, *log*_2_FoldChange=−2.99), Bomanins (BomBc1, *log*_2_FoldChange=−2.05; BomS3, *log*_2_FoldChange=−1.13), and Baramicin (BaraA2, *log*_2_FoldChange=−1.30).

### Lifelong colonized vs. adult recolonized

A focal point of this study is the comparison between generationally colonized flies versus flies only colonized as new adults to give insight into transcriptional changes from germ-free development that cannot be corrected by reintroduction of bacteria. A total of only 29 DEGs, with 25 upregulated and 4 downregulated, were identified between CA and CAA (*log*_2_FoldChange range: −1.04 to 25.12, Table S1). Two histone H2A variants (*His2A:CG33823, His2A:CG33826*) showed the largest fold changes; however, their expression was highly variable across replicates, with reads detected in only a minority of samples and zero counts in other samples. This pattern is inconsistent with genuine biological regulation and likely reflects ambiguous mapping among the highly similar histone H2A paralogous genes (Table S2). These genes were therefore not interpreted biologically.

The remaining differentially expressed genes included mitochondrial Complex I subunits (*mt:ND3*, log_2_FC = 2.23; *mt:ND4L*, log_2_FC = 7.03), adipokinetic hormone (*Akh*, log_2_FC = 1.71), and maltase genes (*Mal-A2, Mal-A4*), consistent with differences in mitochondrial function, energy mobilization, and carbohydrate metabolism between embryonically colonized and adult-recolonized flies.

These findings suggest that flies colonized in early adulthood are nearly equivalent to generationally colonized flies with the exception of a handful of genes that should be studied further, potentially reflecting developmental programming of mitochondrial function (Han et al., 2017) and energy homeostasis by the microbiome (Shin et al., 2011) (Figure 4).

**Figure 4:**
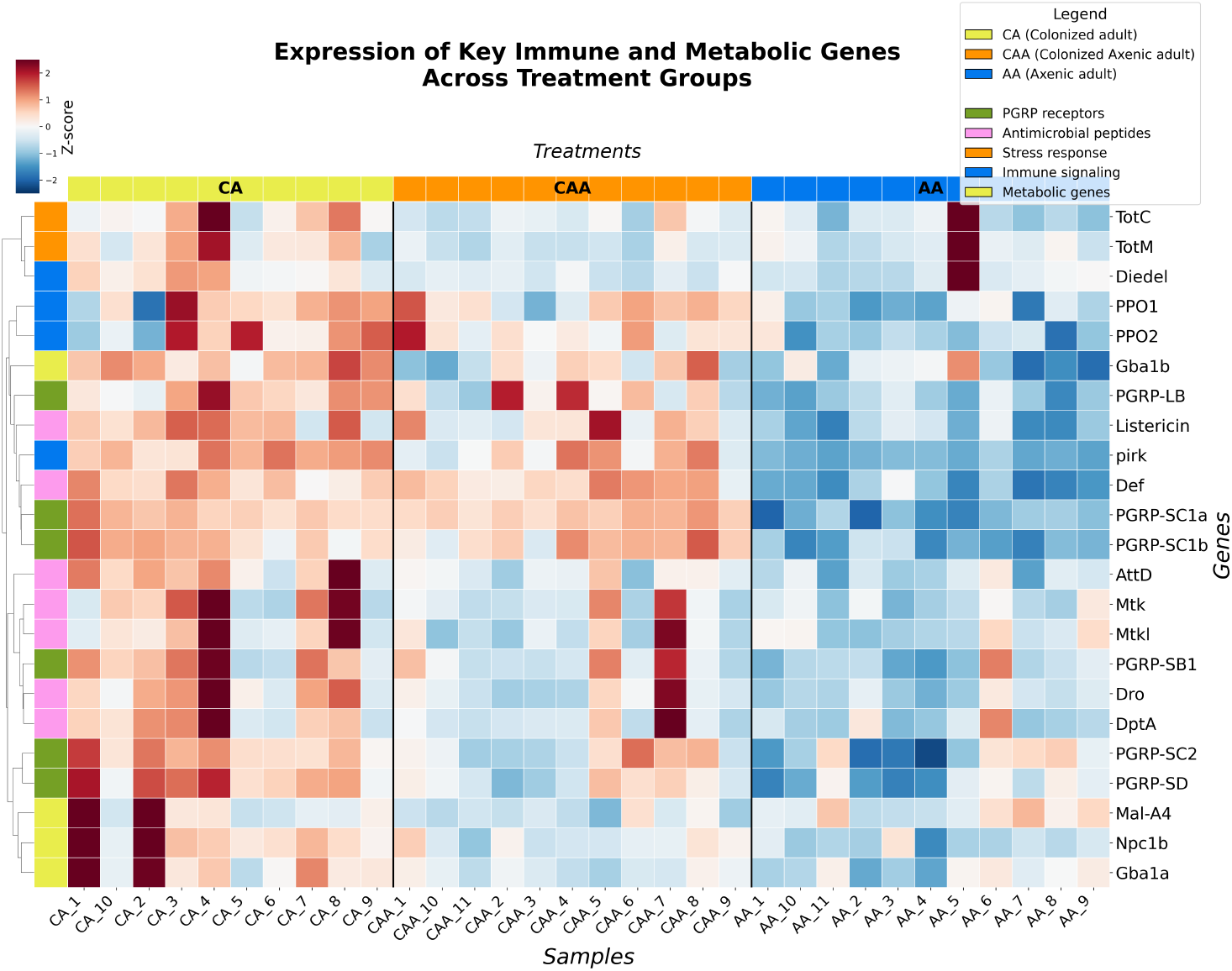
Heatmap of key immune and metabolic gene expression across adult treatment groups. Expression values are Z-score normalized VST counts. Each row represents a specific immune or metabolic gene, and each column represents an individual sample. Gene expression levels are color-coded, with red indicating upregulation and blue indicating downregulation relative to the gene’s mean expression across all samples. Colored bars on the left denote gene categories: PGRP receptors (green), antimicrobial peptides (pink), stress response (orange), immune signaling (blue), and metabolic genes (yellow). Columns are grouped by treatment: CA = Colonized adult (n=10), CAA = Colonized Axenic adult (n=11), AA = Axenic adult (n=11).

**Figure 5:**
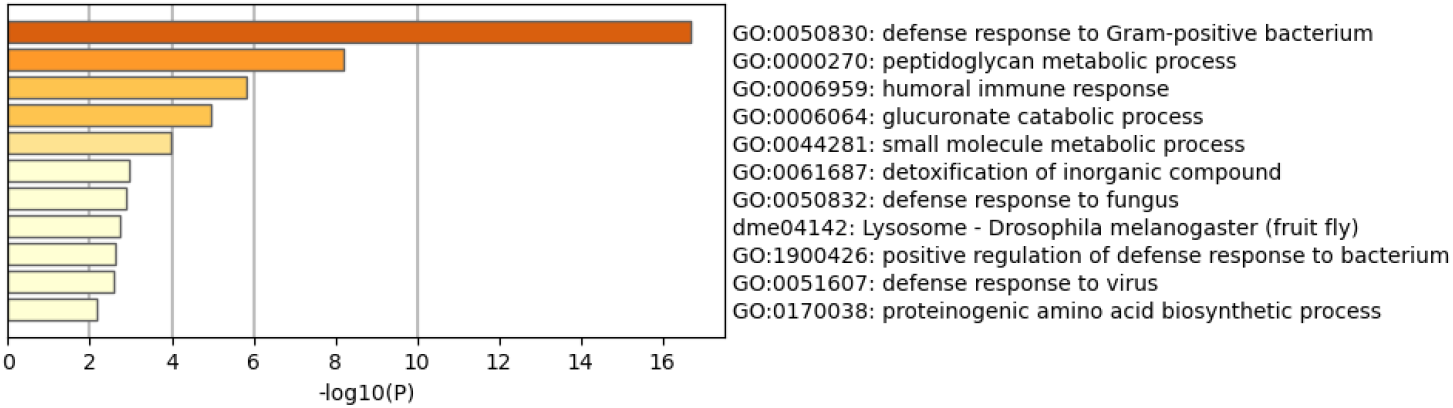
Metascape enrichment heatmap of Gene Ontology terms for genes upregulated in CA vs AA. Top pathways include immune system processes, such as defense response to Gram-positive bacterium.

**Figure 6:**
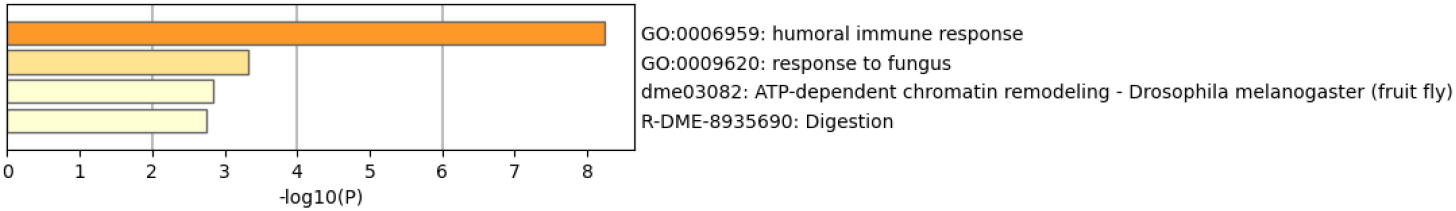
Metascape enrichment heatmap of Gene Ontology terms for genes downregulated in CAA vs AA. Top pathways include humoral immune system processes.

**Figure 7:**
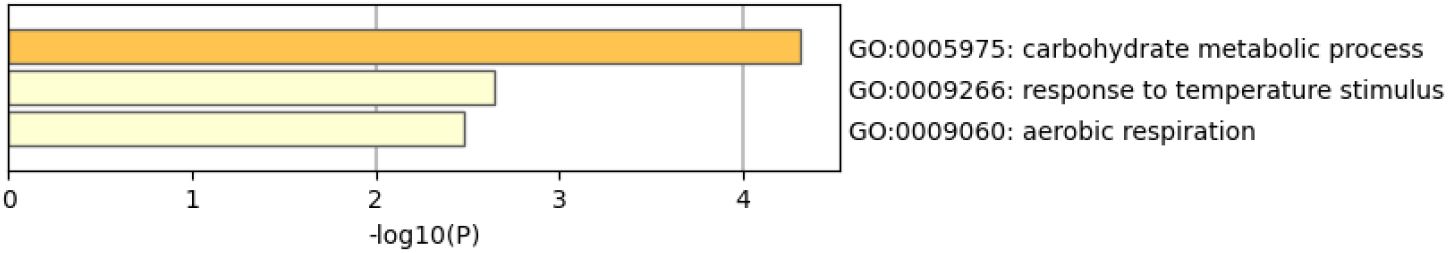
Metascape enrichment heatmap of Gene Ontology terms for genes upregulated in CA vs CAA.

## Discussion

### Mass and Length

The relationship between the gut microbiome and *Drosophila* body size has been well established in developmental contexts. Since 1926, it has been known that flies without a microbiome show normal viability, health, and fecundity when grown on a complete diet of autoclaved yeast (Northrop, 1926). However, when undernourished, axenic larvae have stunted growth, and bacteria rescue the growth deficit (Shin et al., 2011; Storelli et al., 2011). In our study, we used complete nutrition and observed no significant differences in body size between axenic (AA) and colonized (CA) adults, which let us focus on whether other aspects of fly physiology might be affected, such as immunity.

### Transcriptional responses to bacterial colonization

Unsurprisingly, generationally colonized flies showed a robust upregulation of immunity compared to germ-free flies. In contrast, adult colonization of flies that were generationally germ-free upregulated some immune genes but downregulated others. Pattern recognition was activated (PGRP-SC1a/b & PGRP-LB upregulated), but key effector AMPs were suppressed below germ-free baseline levels. The downregulation of Cecropins (CecA2, CecB, CecC; Imd pathway effectors) and Bomanins (BomBc1, BomS3; Toll pathway effectors) below germ-free levels in adult-recolonized flies, rather than simply not induced, suggests active transcriptional repression. This may represent a host strategy to accommodate newly introduced bacteria by reducing antimicrobial pressure, possibly to prevent excessive inflammation against newly encountered microbes that could be beneficial. This suggestion aligns with the hygiene hypothesis framework and the developmental origins of immune function observed in vertebrates (Brodin, 2022). However, reduced AMP expression could compromise defense against bacteria (Loch et al., 2017), potentially explaining why germ-free animals recolonized in adulthood show altered susceptibility to infection, as has been reported for antiviral immunity (Sansone et al., 2015).

The 13 NOT RECOVERED genes were differentially expressed in generationally-colonized flies versus axenic flies and also differentially expressed between generationally-colonized and those colonized only post eclosion. This category included stress-responsive Turandot genes *TotM* (log_2_FC = 1.28) and *TotC* (log_2_FC = 2.21), as well as metabolic genes *Mal-A4, Npc1b*, and *Gba1a*. The *Drosophila* Turandot genes are induced by various stresses, including immune challenge, and contribute to stress tolerance (Ekengren and Hultmark, 2001). Their failure to be mitigated by adult colonization suggests that the stress-adaptive component of the immune response may require early-life bacterial exposure, and this could be a productive area for future studies.

A distinct subset of 10 genes was differentially expressed between CA and CAA but not between either colonized group and axenic controls (CA vs AA or CAA vs AA Figure 8). These genes are sensitive to colonization timing. Two of the genes encode mitochondrial Complex I subunits (mt:ND3 and mt:ND4L); one encodes the neuropeptide Akh, a key regulator of energy mobilization; and CG14075 is annotated as *marmite* (mmt), a putative secreted neuropeptide implicated in nutrient sensing Francisco et al. (2022). *snRNA:U1:21D* and *Mal-A2* may reflect differences in RNA processing and carbohydrate metabolism. Together, these genes suggest that colonization timing may influence pathways related to metabolism and RNA processing. However, because neither colonization state differs detectably from the axenic condition for these genes individually, these results should be interpreted cautiously, with the conservative interpretation being that the timing of colonization has little effect on fly physiology.

**Figure 8:**
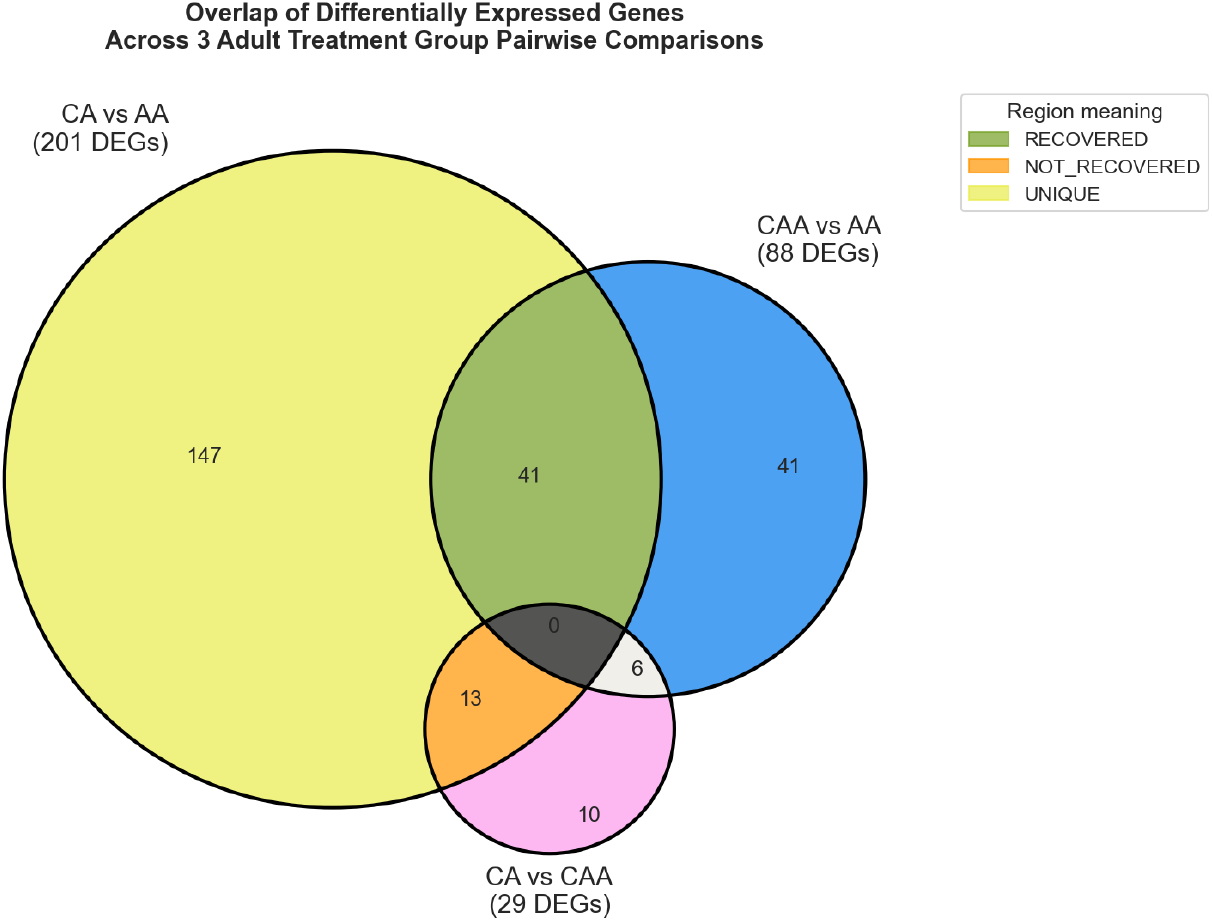
Venn diagram showing overlap of differentially expressed genes across pairwise comparisons of the 3 adult treatment groups. Numbers indicate genes meeting significance criteria (padj ≤ 0.05 | and log_2_ FC | ≥ 0.5) in each comparison. CA = Colonized adult, CAA = Colonized Axenic adult, AA = Axenic adult.

## Conclusions

Our findings reveal remarkable robustness in the host response to the timing of bacterial colonization. Adult recolonization largely restores the gene expression program established by embryonic colonization, with the vast majority of differentially expressed genes, including core immune recognition machinery (PGRPs), pattern recognition receptors, and the negative regulator *pirk*, indistinguishable in early versus late colonization. This suggests that the *Drosophila* immune and metabolic systems retain substantial capacity for adaptation even when colonization is delayed until adulthood.

Nevertheless, a small subset of differentially expressed genes cannot be restored by late colonization, including mitochondrial genes, stress-responsive Turandot genes (TotM, TotC), the immune effector Diedel, and genes involved in metabolism (*Akh, Mal-A4, Npc1b, Gba1a*). These genes provide a focused starting point for future studies investigating how colonization timing influences host physiology. Functional characterization of these candidates (e.g. genetic manipulation or targeted metabolomics) will help determine whether early-life colonization confers lasting physiological advantages, consistent with the developmental origins framework observed in vertebrate host-microbe interactions (Brodin, 2022).

## Limitations of the study

Our study reports the gene expression differences of whole flies, so changes that we observe are due to a combination of changes in the transcriptomes of individual cell types as well as changes in the abundances of individual cell types, which could lead to large differences between treatments. The fact that we see small differences between treatments is consistent with robustness of fly development, as noted by previous studies, e.g. (Elya et al., 2016; Northrop, 1926). Given the few differentially-expressed genes we observed between flies inoculated at birth versus at adulthood (CA and CAA treatments), we feel that even these results should be treated with caution because if a central metabolic regulatory gene like *ahk* were to be stably differentially expressed, we would expect to see widespread physiological changes apparent in the transcriptome of the flies, which we do not.

## Materials and Methods

### Fly Stocks and Maintenance

*Drosophila melanogaster* Canton-S were maintained on standard cornmeal–molasses media (water, molasses, cornmeal, agar, yeast, Tegosept, propionic acid) at 25 ° C in a 12 h light:12 h dark (LD) cycle, with ~65% humidity, as described in (Dodge et al., 2023).

### Bacterial Strains and Culture Conditions

Our bacterial consortium comprises seven strains that act as a commensal in *D. melanogaster*, including *Lactiplantibacillus plantarum* (ZTG301), *Lactobacillus brevis* (ZTG304), *Acetobacter malorum* (ZTG303), *Acetobacter tropicalis* (ZTG310), *Acetobacter orientalis* (ZTG331), *Acetobacter cerevisiae* (ZTG313), and *Acetobacter sicerae* (ZTG360) (Dodge and Ludington, 2023; Dodge et al., 2023; Obadia et al., 2017). Bacterial cultures were prepared and standardized to 10^6^ colony-forming units (CFU) per mL as described (Dodge et al., 2023).

The volume needed was calculated using the formula:

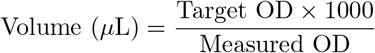

The target OD_600_ values corresponding to 10^6^ CFU/mL for each strain were empirically determined:

### Experimental design

Germ-free (axenic) flies were generated by bleach dechorionation (Kietz et al., 2018; Koyle et al., 2016). Five experimental groups were established.

Embryonic Inoculation: colonized stocks were placed in vials pre-inoculated with 50 µL of the seven-strain bacterial mixture. For colonization, the aforementioned seven species were combined at a total of 3.5 × 10^5^ CFU and applied to the food surface of each vial. Flies developed in the presence of bacteria throughout larval and pupal stages. Newly eclosed (within 24h) P2 adults were collected for analysis (colonized newly eclosed). Following initial collection, a subset of embryonic inoculated adults was transferred to fresh vials containing food inoculated with the same bacterial community. These flies were maintained for an additional 5-7 days post-eclosion before collection (colonized adult).

Axenic controls: Germ-free embryos were reared on sterile food throughout development. Newly eclosed (within 24 h) P2 adults were collected to establish baselines for germ-free physiology (axenic newly eclosed).

Adult Recolonization: After eclosion, a subset of axenic controls was transferred to vials containing food pre-inoculated with the same 50 µL of the seven-strain bacterial mixture as those used for Embryonic Inoculation. Flies were collected after 5–7 days of colonization (colonized axenic adult).

Axenic adult: A separate subset of axenic control was transferred to fresh sterile food for 5–7 days post-eclosion and then collected as long-term germ-free controls.

Colonization status was verified by plating vial contents on MRS and MYPL agar.

### Body Size Measurements

Body length was measured from CO_2_ anesthetized flies imaged under a Zeiss Stemi 508 stereomicroscope. A millimeter ruler placed in the focal plane of each image was used for scale calibration in FIJI/ImageJ (Schindelin et al., 2012). Body length was measured in two segments: head length (anterior to posterior tip of head) and trunk length (posterior tip of head to posterior tip of abdomen) (Figure 9). Body mass was determined by weighing groups of five flies on a microbalance and calculating individual mass.

**Figure 9:**
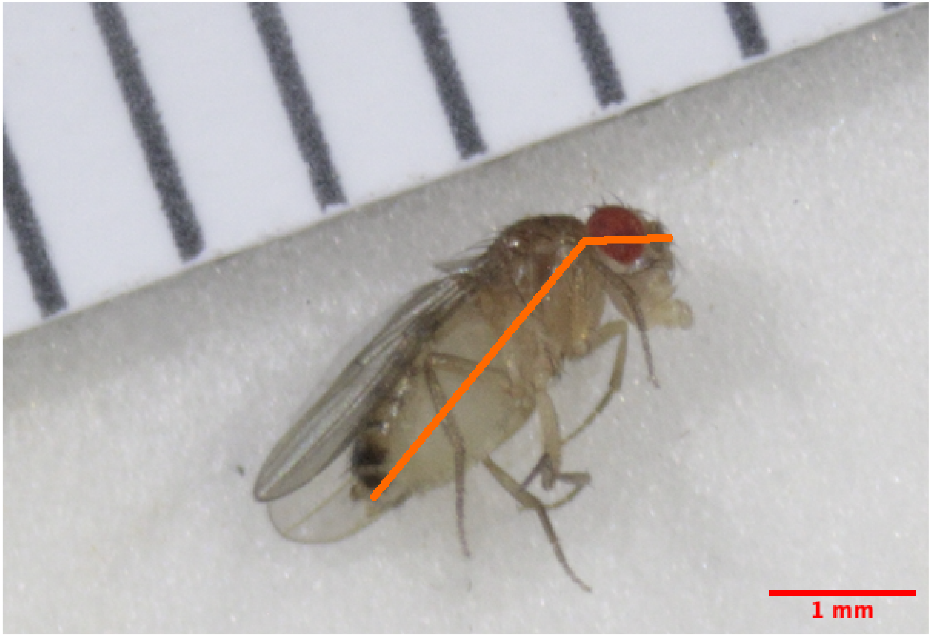
Example of how to measure the length of flies. An image of a female fly with measurements.

To assess treatment effects, we compared body size among adult groups (AA, CAA, and CA). Initial t-test analysis suggested that CAA flies were significantly smaller than CA flies in body length (2.568 ± 0.043 mm vs 2.803 ± 0.042 mm, p = 0.0066 for females). However, given the small sample sizes (n = 3 per group), we applied the Mann-Whitney U test, which does not assume normality and is more appropriate for small samples. Since with n = 3 per group, the minimum achievable two-sided p-value is 0.10, limiting statistical power to detect subtle effects, One-sided Mann-Whitney U tests are also used for analysis. Both U tests revealed no significant differences among treatment groups (all *p* ≥ 0.05).

Despite the lack of statistical significance, due to limited sample size, it remains plausible that CAA flies exhibited smaller body size relative to both AA and CA groups. However, any such effects would appear to be quite small in magnitude.

### RNA Extraction

Total RNA was extracted from pools of five female flies using TRIzol reagent following the total RNA extraction from the Li lab at U Conn (Li Laboratory, 2017). Modifications included the use of Phase Lock Gel Heavy tubes for improved phase separation and bead-beating homogenization of whole flies. RNA quality was assessed using NanoDrop spectrophotometry (A_260_/A_280_ = 1.6–2.1) and Agilent 2100 Bioanalyzer (RIN ≥ 6.5).

RNA samples (one microgram each) were processed using the Illumina TruSeq Stranded mRNA Library Prep kit (Cat #20020594) with IDT for Illumina TruSeq RNA UD Index v2 96 indices (Cat #20040871) according to the manufacturer’s instructions. DNA libraries were quantified on a Qubit v3 and examined on Agilent Bioanalyzer using High Sensitivity DNA reagents. Sequencing was performed on the Illumina NextSeq 1000 platform, generating 100bp single-ended reads with a combined yield of 6.31 to 31.6 million reads per sample. Reads were aligned to the *D. melanogaster* reference genome (BDGP6.54, Ensembl release 115) using the nf-core/rnaseq pipeline (Ewels et al., 2020). One sample with low complexity identified in the MultiQC report was excluded from analysis.

## Data availability

Sequencing data is available at NCBI GEO accession GSE341440: https://www.ncbi.nlm.nih.gov/geo/query/acc.cgi?acc=GSE341440. Analysis code is available on GitHub (https://github.com/exporterdream/Protocol-for-Gut-bacteria-on-general-physiology-exp) and archived in Zenodo (https://doi.org/10.5281/zenodo.21941636).

### Differential expression analysis

Differential expression analysis was performed using DESeq2 (v1.42.0) (Love et al., 2014) in R (version 4.4.0) (Team, 2024). Genes with fewer than 50 total counts across all samples were excluded, yielding 12,482 genes for analysis. A single model containing all three adult groups was fitted, followed by pairwise comparisons to identify differentially expressed genes. Genes with Benjamini-Hochberg adjusted p-value ≤ 0.05 and | log_2_ FC| ≥ 0.5 were considered differentially expressed. Genes with sparse or low baseline counts can exhibit large log_2_FC values whose magnitude is unreliable, and such genes are interpreted with caution.

### Gene classification and pathway analysis

Colonization-responsive genes were classified into three categories based on overlap analysis: RECOVERED genes, NOT RECOVERED genes, and UNIQUE genes (Figure 8). Pathway enrichment analysis was performed using Metascape Zhou et al. (2019) with default parameters.

### Statistical Analysis

Body size comparisons were performed using t-tests and Mann-Whitney U test(Mann and Whitney, 1947; Virtanen and et al., 2020). Statistical significance was set at *p <* 0.05.

## Acknowledgment

We thank Angela Xu (Ludington lab) for training in RNA extraction and bacterial culture techniques. We thank Joseph Tran (Carnegie Institution for Science) for performing RNA sequencing library preparation and sequencing, and Xiaobin Zheng (Carnegie Institution for Science) for assistance with read mapping and data processing.

## Supplementary Material

**Table S1:** 29 DEGs from the CA vs CAA comparison with putative functional annotation and overlap with CA vs AA and CAA vs AA comparisons. Genes labeled “Shared with CA vs AA” were differentially expressed in both CA vs AA and CA vs CAA; genes labeled “Shared with CAA vs AA” were differentially expressed in both CAA vs AA and CA vs CAA; genes labeled “Unique to CA vs CAA” were detected only in CA vs CAA. Log_2_ fold-change values shown are from the CA vs CAA comparison.

| Gene Name | FlyBase ID | log<sub>2</sub>FC | Final Function | Overlap Category |
| --- | --- | --- | --- | --- |
| Ldh | FBgn0001258 | 0.649 | Lactate dehydrogenase; glycolytic enzyme | Shared with CAA vs AA |
| Mal-A2 | FBgn0002569 | 0.521 | Maltase A2; α-glucosidase | Unique to CA vs CAA |
| snRNA:U1:21D | FBgn0003916 | 7.602 | U1 small nuclear RNA; spliceosome component | Unique to CA vs CAA |
| Akh | FBgn0004552 | 1.707 | Adipokinetic hormone; neuropeptide | Unique to CA vs CAA |
| mt:ND3 | FBgn0013681 | 2.226 | Mitochondrial NADH dehydrogenase subunit 3 | Unique to CA vs CAA |
| mt:ND4L | FBgn0013683 | 7.027 | Mitochondrial NADH dehydrogenase subunit 4L | Unique to CA vs CAA |
| mt:srRNA | FBgn0013688 | 3.267 | Mitochondrial small ribosomal RNA | Shared with CAA vs AA |
| CG17260 | FBgn0031498 | 0.760 | Uncharacterized protein | Shared with CAA vs AA |
| TotM | FBgn0031701 | 1.515 | Turandot M; stress response protein | Shared with CA vs AA |
| Mal-A4 | FBgn0033294 | 2.029 | Maltase A4; α-glucosidase | Shared with CA vs AA |
| CG10910 | FBgn0034289 | 0.695 | Uncharacterized protein | Unique to CA vs CAA |
| CG10912 | FBgn0034296 | 0.596 | Uncharacterized protein | Shared with CA vs AA |
| CG9119 | FBgn0035189 | 0.988 | Uncharacterized protein | Shared with CA vs AA |
| CG11594 | FBgn0035484 | 0.661 | Uncharacterized protein | Shared with CA vs AA |
| CG9760 | FBgn0036259 | -1.045 | Uncharacterized protein | Unique to CA vs CAA |
| CG14075 | FBgn0036835 | 1.796 | Putative secreted peptide | Unique to CA vs CAA |
| CG9297 | FBgn0038181 | -0.675 | Uncharacterized protein | Shared with CA vs AA |
| Diedel | FBgn0039666 | 1.276 | Immune-related protein | Shared with CA vs AA |
| CG7567 | FBgn0039670 | -0.557 | Uncharacterized protein | Shared with CA vs AA |
| CG3699 | FBgn0040349 | 1.357 | Uncharacterized protein | Unique to CA vs CAA |
| TotC | FBgn0044812 | 1.681 | Turandot C; stress response protein | Shared with CA vs AA |
| Gba1a | FBgn0051148 | 1.481 | Glucocerebrosidase 1a; sphingolipid metabolism | Shared with CA vs AA |
| lncRNA:CR32773 | FBgn0052773 | -0.628 | Long non-coding RNA | Shared with CA vs AA |
| His2A:CG33823 | FBgn0053823 | 25.121 | Histone H2A variant; chromatin-associated | Shared with CAA vs AA |
| His2A:CG33826 | FBgn0053826 | 23.786 | Histone H2A variant; chromatin-associated | Shared with CAA vs AA |
| CG42335 | FBgn0259237 | 0.890 | Uncharacterized protein | Shared with CA vs AA |
| Npc1b | FBgn0261675 | 1.177 | Cholesterol trafficking protein | Shared with CA vs AA |
| BomT1 | FBgn0262838 | 1.978 | Bomanin T1; antimicrobial peptide | Shared with CAA vs AA |
| roo{ }339 | FBti0019148 | 1.468 | Transposable element | Unique to CA vs CAA |

**Table S2:** Genes with unusually large log_2_ fold-change estimates are shown with their total raw counts and number of samples with nonzero expression across 57 libraries. All fold-change comparisons are between adult groups (CA, CAA, AA). Although those genes passed the count filter, their signal was concentrated in only a few samples, suggesting ambiguous mapping among highly similar paralogous or repetitive genes rather than genuine biological regulation.

| Gene | $\vert \log_2 \text{FC}\vert $ | Total raw counts | Nonzero samples |
| --- | --- | --- | --- |
| His2A:CG33826 | 23.79 (CA vs CAA), 25.21 (CAA vs AA) | 2,544 | 6/57 |
| His2A:CG33823 | 25.12 (CA vs CAA), 25.08 (CAA vs AA) | 2,724 | 6/57 |
| Juan{}257 | 20.85 (CA vs AA), 22.87 (CAA vs AA) | 805 | 6/57 |
| His2A:CG33820 | 22.72 (CAA vs AA) | 2,317 | 5/57 |
| His4:CG33893 | 21.95 (CA vs AA), 22.29 (CAA vs AA) | 875 | 6/57 |
| 5.8SrRNA:CR45852 | 22.15 (CA vs AA), 21.40 (CAA vs AA) | 22,093 | 15/57 |
| tal-1A | 20.15 (CAA vs AA) | 2,878 | 9/57 |
| His4:CG33897 | 9.54 (CA vs AA) | 1,106 | 9/57 |
| snRNA:U1:21D | 7.60 (CA vs CAA) | 1,091 | 18/57 |
| mt:ND4L | 7.03 (CA vs CAA) | 1,175 | 32/57 |
| invader1{}1301 | 3.63 (CA vs AA), 6.59 (CAA vs AA) | 1,925 | 27/57 |
| CG31131 | 5.34 (CAA vs AA) | 218 | 24/57 |

**Figure S1:**
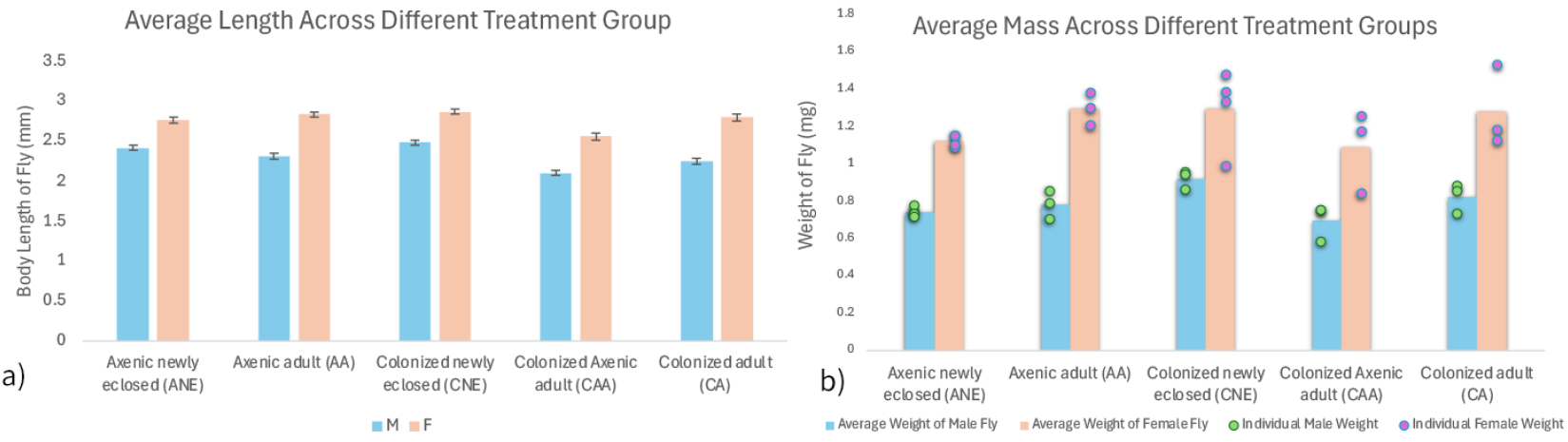
Comparison between five condition. a) Comparison between length of different sex, with standard error as error bar. b) Comparison of weight between sex.

**Figure S2:**
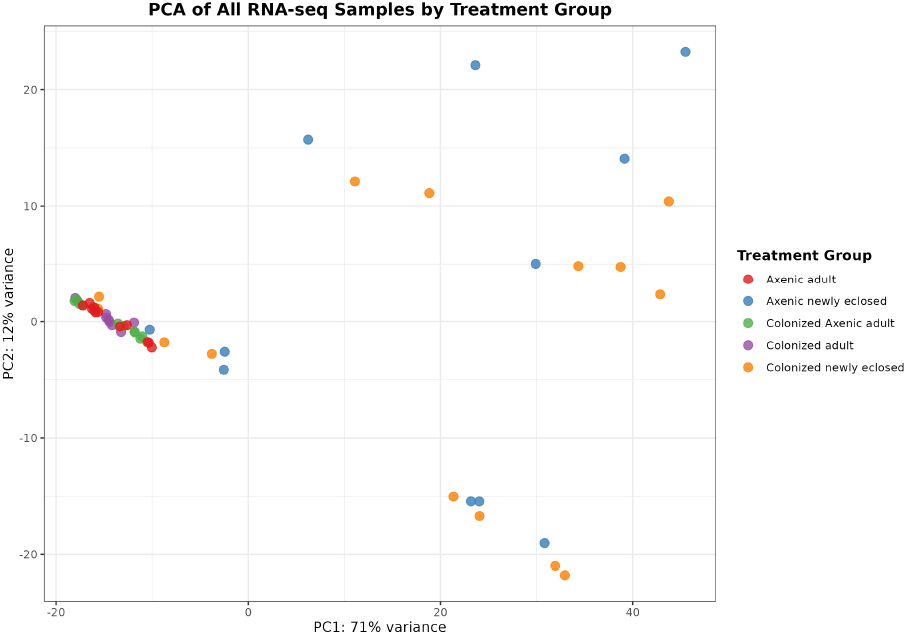
Principal Component Analysis (PCA) of RNA-seq samples. Colors indicate treatment groups as defined in the legend. Clustering visualization of all samples across all treatment groups.

## Notes

https://www.ncbi.nlm.nih.gov/geo/query/acc.cgi?acc=GSE341440

https://github.com/exporterdream/Protocol-for-Gut-bacteria-on-general-physiology-exp

https://doi.org/10.5281/zenodo.21941636

### Competing Interest Statement

The authors have declared no competing interest.

### Summary of Updates

Figure 3 labels were added to panels A-C and Table S2 was added. Minor text edits were incorporated.

### Funding

- U.S. National Science Foundation: 2549672

